# Transcript-specific enrichment enables profiling rare cell states via scRNA-seq

**DOI:** 10.1101/2024.03.27.587039

**Authors:** Tsion Abay, Robert R. Stickels, Meril T. Takizawa, Benan N. Nalbant, Yu-Hsin Hsieh, Sidney Hwang, Catherine Snopkowski, Kenny Kwok Hei Yu, Zaki Abou-Mrad, Viviane Tabar, Leif S. Ludwig, Ronan Chaligné, Ansuman T. Satpathy, Caleb A. Lareau

**Affiliations:** Gladstone-UCSF Institute of Genomic Immunology, San Francisco, CA, USA; Department of Pathology, Stanford University, Stanford CA, USA; Program in Biological and Biomedical Sciences, Harvard University, Boston, MA, USA; Single-cell Analytics Innovation Lab, Memorial Sloan Kettering Cancer Center, New York, NY, USA; Computational and Systems Biology Program, Memorial Sloan Kettering Cancer Center, New York, NY, USA; Charité Universitätsmedizin Berlin, Germany; Berlin Institute of Health at Charité – Universitätsmedizin Berlin, Berlin, Germany; Max-Delbrück-Center for Molecular Medicine in the Helmholtz Association (MDC), Berlin Institute for Medical Systems Biology (BIMSB), Berlin, Germany; Department of Neurosurgery, Memorial Sloan Kettering Cancer Center, New York, NY, USA; Parker Institute for Cancer Immunotherapy, San Francisco, CA, USA

**Author notes:** These authors contributed equally (Tsion Abay, Robert R. Stickels, Meril T. Takizawa).

## Abstract

Single-cell genomics technologies have accelerated our understanding of cell-state heterogeneity in diverse contexts. Although single-cell RNA sequencing (scRNA-seq) identifies many rare populations of interest that express specific marker transcript combinations, traditional flow sorting limits our ability to enrich these populations for further profiling, including requiring cell surface markers with high-fidelity antibodies. Additionally, many single-cell studies require the isolation of nuclei from tissue, eliminating the ability to enrich learned rare cell states based on extranuclear protein markers. To address these limitations, we describe Programmable Enrichment via RNA Flow-FISH by sequencing (PERFF-seq), a scalable assay that enables scRNA-seq profiling of subpopulations from complex cellular mixtures defined by the presence or absence of specific RNA transcripts. Across immune populations (*n* = 141,227 cells) and fresh-frozen and formalin-fixed paraffin-embedded brain tissue (*n* = 29,522 nuclei), we demonstrate the sorting logic that can be used to enrich for cell populations via RNA-based cytometry followed by high-throughput scRNA-seq. Our approach provides a rational, programmable method for studying rare populations identified by one or more marker transcripts.

## Introduction

Rare cell types are found across diverse biological contexts and often participate critically in tissue function, represent key transitional states, or define incipient disease. Single-cell RNA-seq has powered the discovery of many minority populations that can be defined by cell-type-specific transcript combinations, but their limited sampling makes them difficult to study. For example, *CFTR*^+^ pulmonary ionocytes likely mediate the pathogenesis of cystic fibrosis, but make up only 1 in 200 human lung epithelial cells^1^. Similarly, enteric neurons present at a frequency of 1 in 300 in the colon^2^ and *AXL*^+^ *SIGLEC6*^+^ dendritic cells (AS DCs) at 1 in 5,000 peripheral blood mononuclear cells (PBMCs)^3,4^. More recently, we identified a rare population of chimeric antigen receptor (CAR) T cells that reactivate human herpesvirus 6 at a frequency of ∼1 in 10,000 cells in infusion products^5^, which may contribute to the etiology of HHV-6 encephalitis in patients receiving cell therapies. Though these anecdotes represent diverse populations and tissue types, the conceptual mode of discovery for these rare populations has been consistent: genomic profiles of ∼10^5^-10^7^ cells were generated, yielding ∼10^1^-10^3^ events of interest. The tremendous resources required to define very rare but consequential populations of cells limit the power for downstream analyses, including the identification of transcriptional heterogeneity within these populations, inference of additional marker genes, and analysis of gene regulatory networks.

After these populations are identified and defined by marker transcripts via scRNA-seq, challenges persist in isolating them for further characterization. A frequently used approach is the enrichment or depletion of cells expressing specific surface proteins via fluorescence-activated cell sorting (FACS). However, as these rare populations are defined through transcriptomics analyses, analogous surface proteins may not be defined or high-quality antibodies available. Further, many archived sample preparations require nuclei dissociation steps from frozen or formalin-fixed paraffin-embedded (FFPE) tissue samples, which eliminates the possibility of enriching or depleting based on any non-nuclear protein. Though the profiling of intranuclear proteins and scRNA-seq was recently demonstrated^6^, high-quality antibodies recognizing transcription factors can be inaccessible due to a lack of highly structured antigens available for targeting^7^. These limitations motivate an approach that enriches for either cells or nuclei based on individual or arbitrary combinations of RNA markers upstream of additional scRNA-seq profiling.

To this end, we report the development of Programmable Enrichment via RNA Flow-FISH by sequencing (PERFF-seq). Our assay utilizes fluorescence *in situ* hybridization (FISH) to specifically label RNA(s) that enrich populations of interest and is compatible with downstream high-quality transcriptional profiling via high-throughput droplet-based scRNA-seq. We identify parameters in currently utilized FISH workflows that diminish downstream scRNA-seq performance and determine conditions that yield high-quality data on par with libraries derived from commercially available protocols. We demonstrate the broad applicability of PERFF-seq to enrich immune cell subsets using one or more RNA markers, including the mRNA of transcription factors. Further, we show the compatibility of our protocol with nuclei extracted from frozen or FFPE tissue samples. Together, PERFF-seq enables the efficient enrichment and high-throughput profiling of cells and nuclei populations of interest at high throughput, using logic-gated sorting across heterogeneous cell and tissue types.

## Results

### Assay rationale and overview

Although tissues preserved in formaldehyde, including FFPE tissue blocks, are broadly available, extracting RNA from these sources has remained a major challenge^8^. Notably, formaldehyde-associated RNA degradation is a major limiting factor in generating high-quality libraries from fixed cells, as the degraded RNA molecules cannot be reliably reverse-transcribed for conventional downstream analyses. A recently introduced workflow from 10x Genomics, termed Single Cell Gene Expression Flex^9^ (‘Flex’ hereafter), facilitates the profiling of fixed cells using whole transcriptome probe pairs, by detecting and quantifying transcripts based on the adjacent hybridization and subsequent ligation of probes in droplets (**Extended Data Fig. 1a**). In parallel, advances in FISH technologies, including hybridization chain reaction (HCR), generate an amplified fluorophore signal with high signal-to-noise separation^10^ upon hybridization to RNA molecules of interest^11,12^, enabling flow cytometry-based detection and enrichment of specific populations at very high throughput. Notably, all RNA FISH technologies require an irreversible fixation step, which makes it impossible to apply conventional reverse transcription-based scRNA-seq technologies. We thus envisioned a programmable method (i.e., with user-defined target genes) for enriching cells via RNA FISH and flow cytometry-based sorting, followed by single-cell transcriptome profiling via the Flex scRNA-seq workflow (**Fig. 1a**). Such an approach would enable the study of heterogeneity underlying cell states based on the abundance of any marker genes, including the rare cell states noted above^123,45^.

**Figure 1.**
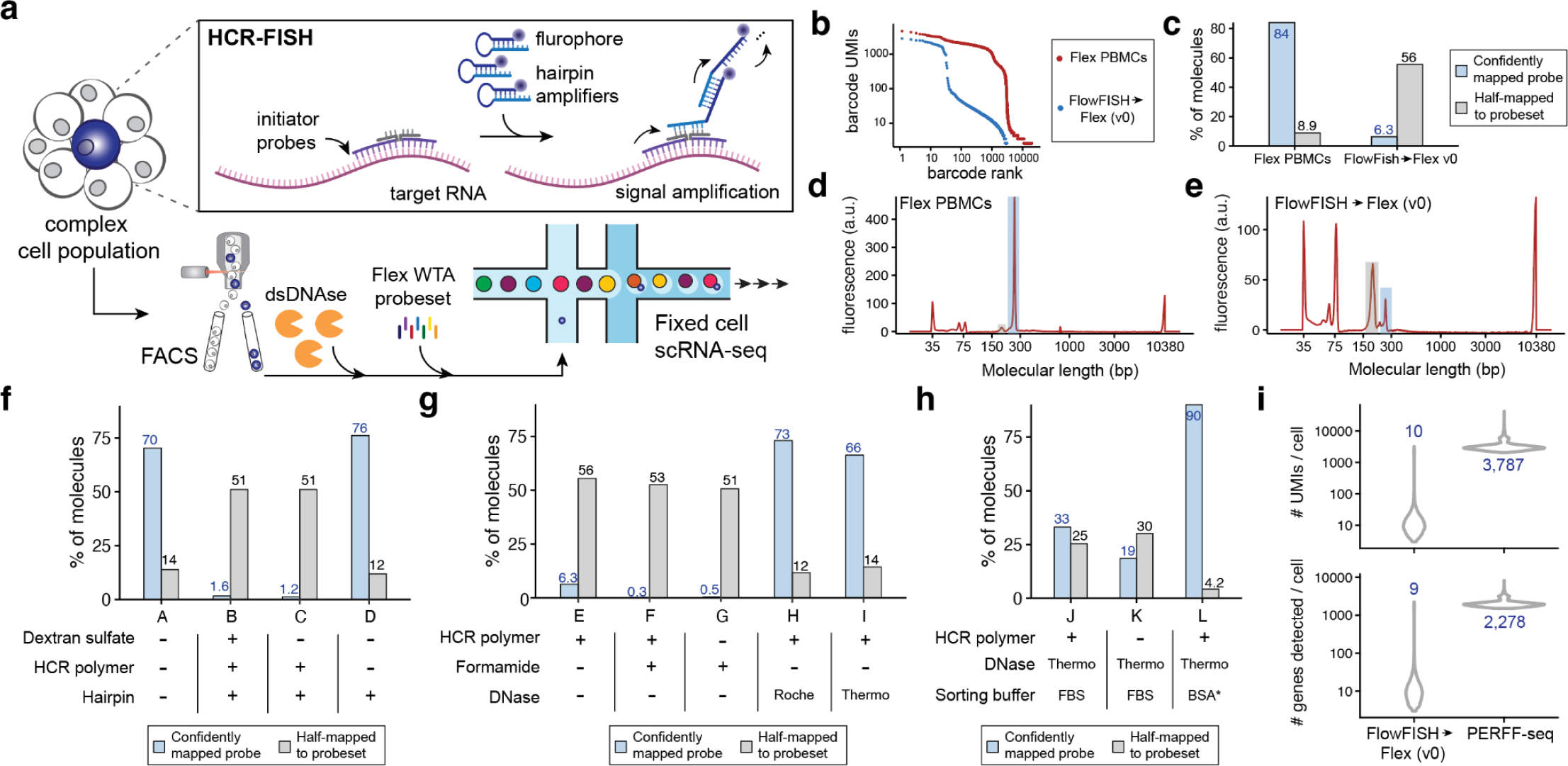
Rationale and development of PERFF-seq. **(a)** Schematic of the PERFF-seq assay. Target RNA(s) are bound by pairs of adjacent initiator probes that ensure specificity. Hairpin amplifiers unzip and hybridize iteratively to generate strong fluorescent signal and enable FACS prior to single-cell profiling with the droplet-based scRNA-seq Flex kit. **(b)** Knee plot of cells profiled with standard Flex versus HCR-FlowFISH sorted cells. **(c)** Fraction of reads fully mapping (blue) or half-mapping (grey) to the reference probe set. **(d)** Bioanalyzer traces highlighting expected product size of the full probe (∼260 bp; blue) and half probe (∼190 bp; grey) for a high-quality Flex library. **(e)** Same as (d) but for the FlowFISH → Flex v0 experiment. **(f)** Experiments identifying the HCR polymer as the corrupting agent for data quality. **(g)** Conditions screened for polymer stripping, including DNase and formamide treatments. **(h)** Sorting buffer components analyzed to improve data quality. **(i)** UMIs (top) and genes (bottom) detected per cell comparing initial FlowFISH → Flex v0 experiment, from (b) to the final PERFF-seq library, L, from panel (h). Values plotted in **c** and **f–i** represent overall library values for a single replicate.

### PERFF-seq development and quality control

We first tested the simple concatenation of the HCR-FlowFISH workflow^11,13^ with Flex library preparation steps. After sorting PBMCs for the B-cell-specific marker gene *MS4A1* (encoding CD20), we captured cells and compared quality control measures to a standard Flex library of only PBMCs (**Extended Data Table 1; Methods**). Despite targeting a similar number of cells (∼3,000), the FlowFISH-sorted cells yielded far fewer cells with overall worse per-cell data quality than the standard Flex library (**Fig. 1b; Extended Data Table 2**).

We noticed that this FlowFISH→Flex v0 library contained a high number of reads half-mapping to the probeset, indicating poor ligation efficiency of Flex gene probe pairs, which is likely due to compromised specific hybridization (**Fig. 1c; Extended Data Fig. 1b**). Specifically, among reads captured in the library, we observed a 15-fold increase of expected probe hybridization sequences containing the probe capture sequence, reflecting the barcoding of unligated product (**Extended Data Fig. 1c**; **Methods**). These inferences were corroborated by Bioanalyzer traces for these two libraries, which verified that the extension of the truncated probes severely limited per-cell data quality, consistent with an application note from 10x Genomics (**Fig. 1d,e; Methods**). Thus, to our surprise, the direct integration of FlowFISH into fixed-cell scRNA-seq is not compatible with attaining high-quality genomics data.

We next interrogated the impact of varying individual FISH workflow components, including dextran sulfate buffer, which is used for background suppression but can negatively impact enzymatic activity^11,14,15^, and reagents for generating fluorescent signal, including the probe and hairpin combination (HCR-polymer) or the hairpin alone (**Fig. 1f**). Dextran sulfate had no impact, but the generation of the HCR polymer was detrimental to data quality (1.2% vs 76% full probe reads; **Fig. 1f**). As the formation of the gene probe-hairpin polymer is essential for FISH signal amplification, we reasoned that removing the tethered probe-hairpin polymer after sorting but before scRNA-seq library preparation could rescue transcriptional profiling.

To assess this, we sought to disassemble the HCR polymer via formamide^11,14^ and enzymatic^10^ stripping, both of which have been used in imaging and microscopy analyses (**Methods**). Though formamide enabled probe-hairpin polymer stripping as expected (**Extended Data Fig. 1d)**, it inhibited the capture of virtually any ligation product (0.5% reads mapping to the full probe set), irrespective of the presence of HCR-polymer (**Fig. 1g**). Alternatively, digestion of the HCR polymer via dsDNase (Thermo) reduced the fluorescence signal as expected, but yielded markedly higher library complexity and ligation efficiency, on par with our original Flex libraries (**Fig. 1g; Extended Data Fig. 1e**). Sorting using BSA with an RNase inhibitor (rather than FBS as used by prior HCR-FlowFISH applications^13^) further improved scRNA-seq library quality (**Fig. 1h; Methods**). The result was a FlowFISH-sorted Flex library with 90% of molecules mapping to a full probe set and 4.2% mapping to half-ligated probes, which was confirmed via Bioanalyzer traces (**Extended Data Fig. 1f**) and consistent with our initial Flex library (**Fig. 1c**).

Performing the full workflow with all modifications except the dsDNase step (**Methods**) results in a high fraction of incomplete molecules, demonstrating that polymer stripping is the critical step for avoiding incomplete probe ligation and generating high-quality scRNA-seq libraries (**Extended Data Fig. 1g**). These optimization steps for FlowFISH sorting and profiling via scRNA-seq together comprise the PERFF-seq assay.

### PERFF-seq benchmarking

To assess feasibility, sensitivity, and specificity, we first benchmarked how well HCR-FlowFISH probes enrich for well-defined populations. Staining PBMCs for *ACTB* with two different amplifiers allowed us to assess HCR FISH efficiency on both the 647 and 488 FACS channels. In either channel, 97% of cells were positive for *ACTB*, indicating the high sensitivity of HCR-FlowFISH and compatibility with multiple fluorophores (**Fig. 2a**). To verify sensitivity, we mixed Raji and K562 human cell lines in varying proportions that differ up to two orders of magnitude (**Methods**) we designed probes against *XIST*, a long-noncoding RNA (lncRNA) expressed in cells with XX sex chromosomes (K562), and absent from XY cells (Raji). Flow cytometry confirmed the sensitive and specific detection of *XIST*^+^ populations in as few as 1 in 500 cells (**Fig. 2b**), confirming feasibility with non-coding transcripts, and suggesting that PERFF-seq can be used to study a broad range of RNA molecules.

**Figure 2.**
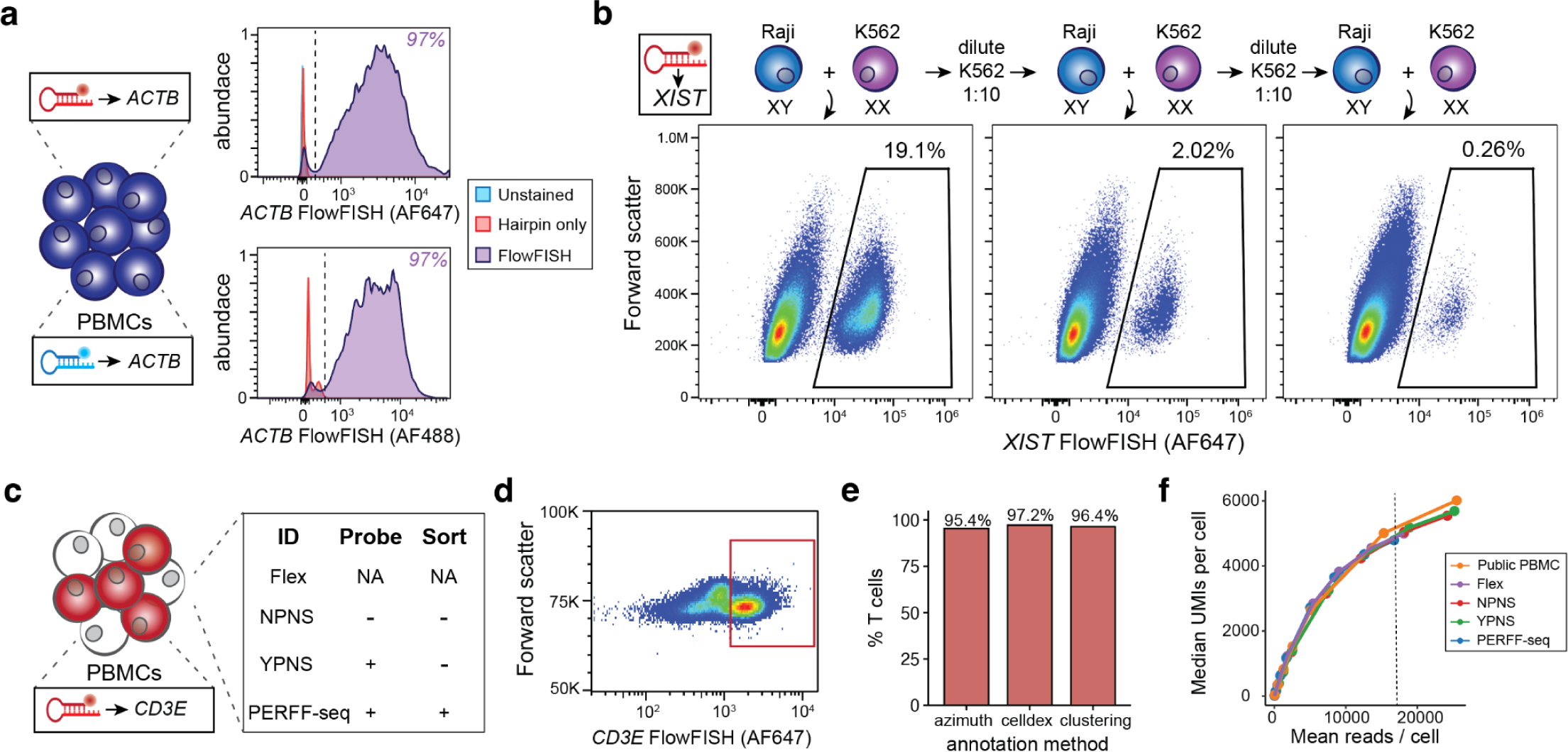
Benchmarking of PERFF-seq. **(a)** Sorting of PMBCs stained with AF647 and AF488-labeled *ACTB* probes. **(b)** Benchmarking of lncRNA *XIST* FlowFISH in serial dilution of K562s (XX / *XIST^+^*) into Rajis (XY / *XIST*^-^). **(c)** PERFF-seq benchmarking experiment for four libraries, including the standard Flex workflow with or without PERFF-seq probe staining or sorting steps. **(d)** Sorting strategy for *CD3E*+ cells for the PERFF-seq library. **(e)** Proportion of cells from the sorted PERFF-seq library annotated as T cells using three different computational methods for classification. **(f)** Downsampling analysis for library saturation and UMI benchmarking. Dotted line represents the mean reads per cell for a final comparison (depth of lowest sample).

To assess the performance of the scRNA-seq profiles, we multiplexed four libraries to compare PERFF-seq differences from the Flex protocol, including probe staining/stripping and sorting steps (**Fig. 2c; Methods**). Across this comparison, we profiled 59,313 cells, recovering the major cell types expected from PBMCs (**Extended Data Fig. 2a,b**). For the PERFF-seq library, we sorted for *CD3E*^+^ cells, yielding libraries with ∼95-97% T cells (**Fig. 2d,e**; **Extended Data Fig. 2c; Methods**). Qualitatively, we did not observe major unexpected population shifts from these four conditions, indicating that PERFF-seq library prep does not meaningfully bias transcript detection relative to the standard Flex workflow. However, we observed that the addition of *CD3E* probes reduced the transcript expression (∼2x reduction of log_2_ counts per million) and percentage of cells (∼24%) detected among T cell populations, relative to *CD3D* (**Extended Data Fig. 2d,e; Methods**). These results suggest that the transcripts targeted by FlowFISH can still be detected via scRNA-seq but warrants caution for interpreting quantitative gene expression levels of the target transcript. Notably, we observed only two differentially expressed genes compared to the standard Flex library, indicating that the additional PERFF-seq steps have minimal impact on whole transcriptome measurements (**Extended Data Fig. 2f; Methods**).

We further compared PERFF-seq results to a gold-standard public PBMC dataset from 10x Genomics. After downsampling for consistency (to 16,750 reads per cell based on the lowest library), we observed almost no loss in data quality; key data quality metrics per cell did not differ by more than ∼10% comparing public Flex (median 5,200 UMIs and median 2,857 genes detected per cell), to our Flex (median 4,885 UMIs and median 2,536 genes), and PERFF-seq libraries (median 4,789 UMIs and median 2,535 genes; **Fig. 2f, Extended Data Table 3; Methods**). Our comprehensive benchmarking analyses together verify that PERFF-seq can be used for cell type enrichment and high-quality scRNA-seq profiling.

### Multi-color programmable enrichment with PERFF-seq

We further benchmarked PERFF-seq by designing multiple probes against three well-described genes in different immune populations in PBMCs, *CD3E*, *MS4A1* (CD20), and *CD4* (**Fig. 3a**). We recovered four cell populations using PERFF-seq with one fluorophore per gene, including a *CD4* and *CD3E* double-positive population that together specifically enrich for CD4^+^ T cells (**Fig. 3b**). scRNA-seq profiling of the four populations in a single reaction using in-line probe-barcoded multiplexing native to Flex yielded a total of 35,220 cells (**Fig. 3c; Methods**). To verify the quality of enrichments, we annotated cell types via three independent methods, showing that PERFF-seq has 70-88% accuracy in recovering the desired cell type labels (**Fig. 3d; Methods**). Analyses of standard PBMC marker genes confirmed the expected populations and recovery of the transcripts used for FlowFISH enrichment and sorting, noting the overall desired depletion of *CD4*^+^*,CD3E^-^*myeloid cells (**Fig. 3e; Extended Data Fig. 3a**).

**Figure 3.**
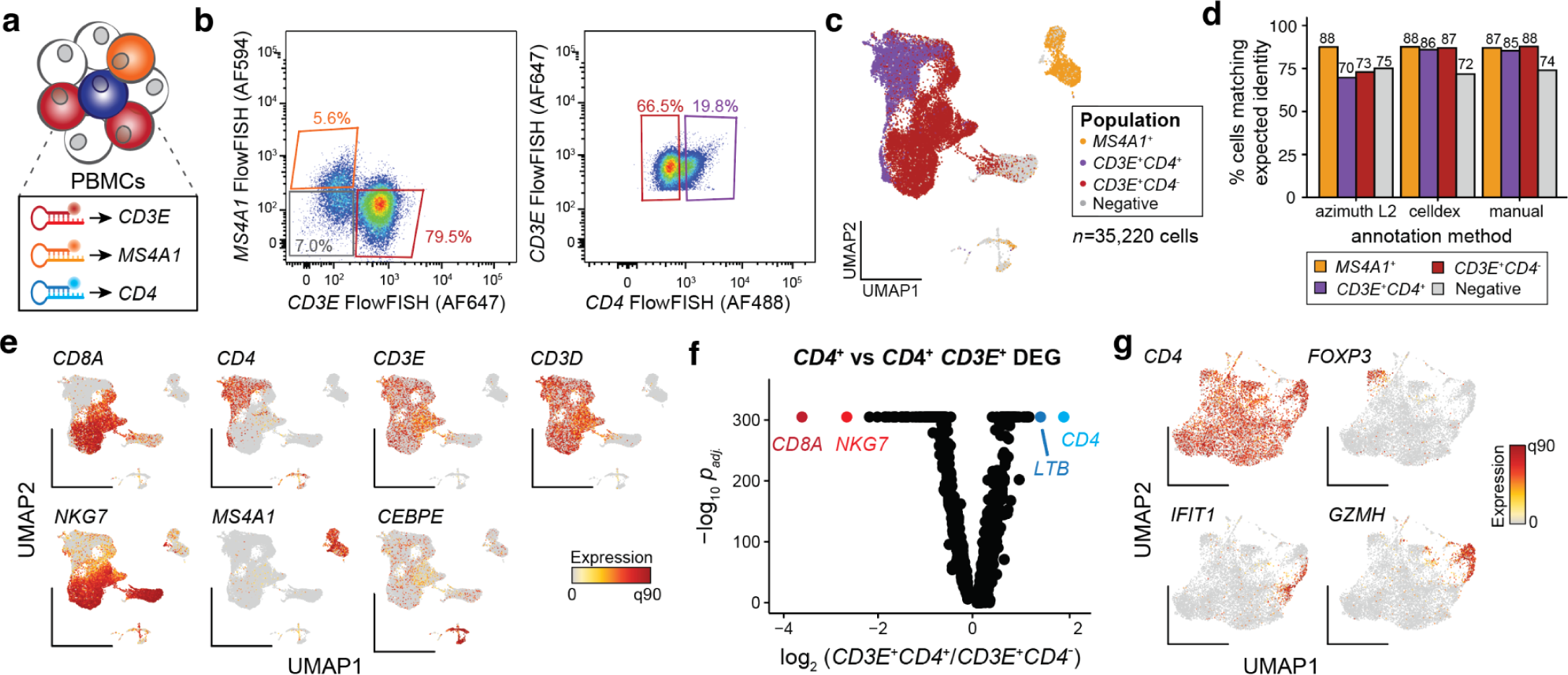
Enrichment of cells via combinatorial logic. **(a)** Schematic of experimental design. Probes targeting three indicated genes, each labeled with distinct fluorophores stain specific populations in PBMCs from a healthy donor. **(b)** FlowFISH signal and sort gates. Percentages represent the overall fraction of events sorted in each gate. **(c)** Reduced dimensionality representation of four populations profiled with PERFF-seq. Colors represent gates drawn from the FlowFISH sort in (b). **(d)** Percent of high-quality cells from PERFF-seq assigned to expected cell types using three distinct annotation methods. Colors represent gates drawn from the FlowFISH sort in (b). **(e)** Annotation of relevant marker genes for populations in redacted dimensionality space, including genes used in the FlowFISH panel. **(f)** Differential gene expression analysis comparing *CD4^+^*and *CD4*^-^ populations from the *CD3E*^+^ sort (panel b, right). Genes corroborating annotation are highlighted. **(g)** Subclustering of *CD3E^+^*/*CD4^+^*cells, highlighting rare subclusters marked by relevant genes.

To confirm the efficacy of the double positive enrichment, we performed differential gene expression analyses of *CD3E*^+^ cells that either co-expressed or were negative for the *CD4* transcript. Reassuringly, the top differential transcripts coincided with key CD4^+^ and CD8^+^ T cell markers (**Fig. 3f; Methods**). Moreover, reclustering the 9,035 cells from the *CD4*,*CD3E* double-positive population, identified major CD4^+^ T cell subsets, including regulatory T cells (T regs) expressing *FOXP3*, naive T cells marked by *LEF1*, interferon-responsive cells expressing *IFIT* genes^16^, and cytotoxic CD4^+^ cells expressing *GZMH* (**Fig. 3g; Extended Data Fig. 3b**). Together, these analyses validate the ability of PERFF-seq to enrich for cellular populations based on specific RNA marker combinations.

### Rare cell states enriched via transcriptional regulators

To expand the study of transcriptional heterogeneity beyond surface protein-defined cell populations, we explored using RNA features typically inaccessible to antibody-based flow cytometry. We elected to enrich populations expressing well-described hematopoietic lineage-defining transcription factors (TFs) *BCL11A* and *SPI1* (encoding PU.1)^17^ which exhibit robust expression across various cell types in public scRNA-seq atlases, and profiled a total of 37,566 cells from *SPI1*^+^*, BCL11A*^+^, and double-negative libraries (**Fig. 4b; Extended Data Fig. 4a,b**). Cell states segregated readily based on the sorted TFs, but also as a function of well-established (surface) marker transcripts (**Fig. 4c; Extended Data Fig. 4c**). Specifically, we observed robust enrichment as 96.0% and 91.1% of cells sorted for *SPI1* and *BCL11A* had non-zero UMI counts for those genes (compared to 13.3% and 9.5% in the unsorted fraction, respectively) with over half of the positive cells having at least 10 UMIs in the enriched libraries (**Fig. 4d, Extended Data Fig. 4d**). Standard clustering and cell type annotation showed a clear skewing of cell states in either enriched TF library, including the consistent and desired depletion of T cells that do not express either TF (**Fig. 4e,f**).

**Figure 4.**
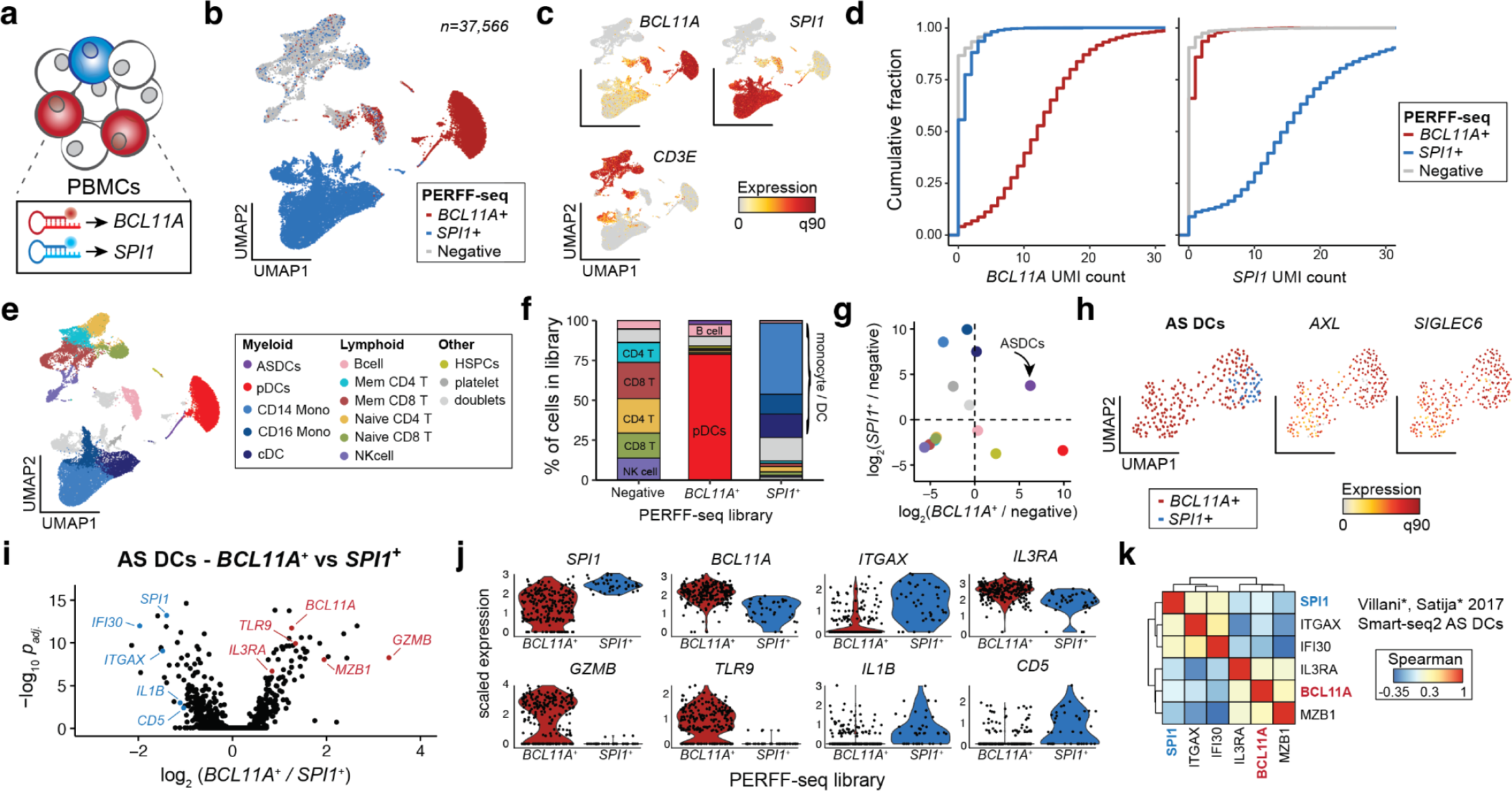
Rare cell states enriched via transcriptional regulators. **(a)** Schematic of human PBMC staining with probes targeting *BCL11A* and *SPI1*. **(b)** UMAP embedding of PERFF-seq profiles from three populations based on TF FlowFISH sorting logic. **(c)** Depiction of marker gene expression across all PERFF-seq profiled cells. **(d)** Empirical cumulative distribution plot of raw UMI counts for *BCL11A* (left) and *SPI1* (right) stratified by the captured PERFF-seq library. **(e)** Annotated cell states from PERFF-seq profiling. **(f)** Proportions of each cell type per library with major cell types labeled. Colors match (e). **(g)** Relative enrichment of each cell type in either the *BCL11A^+^* sort (x-axis) or *SPI1^+^* sort (y-axis) relative to the negative population. *AXL^+^*, *SIGLEC6^+^* dendritic cells (AS DCs) are highlighted as the only enriched population in both sorted populations. Colors as in (e). **(h)** UMAP of AS DCs, highlighting the TF FlowFISH library and defining marker gene expression. **(i)** Volcano plot comparing differentially expressed genes from the two FlowFISH sorted populations. Notable marker genes are highlighted, including known and newly characterized marker genes for AS DC subsets. **(j)** Violin plots of marker genes, stratified by the FlowFISH library. All genes were significantly differentially expressed at a false discovery rate (FDR) <0.01. **(k)** Gene-gene correlations of all AS DCs using the original dataset, highlighting the co-occurrence of TFs from our analysis with established marker genes for the AS DC subsets.

Though *BCL11A* is an essential regulator of B lymphoid development, nearly 75% of the enriched cell population from its PERFF-seq library were pDCs. This enrichment was consistent with bulk^18^ and single-cell expression data^19^ indicating that pDCs express *BCL11A* at 1-2 orders of magnitude higher levels than B cells (**Extended Data. Fig. 4a,e; Methods**). The B cells from the *BCL11A*^+^ PERFF-seq library displayed higher *BCL11A* expression and target gene module scores than B cells from the other two libraries (**Extended Data Fig. 4f; Methods**), indicating that PERFF-seq can enrich for cells with higher TF activities within specific cell types. Further, the *SPI1* staining primarily enriched monocyte and classical dendritic cells (cDCs), consistent with its expression in PBMCs (**Fig. 4f; Extended Data Fig. 4a**).

Next, we asked whether any population was enriched in both TF^+^ populations profiled with PERFF-seq, and indeed, observed that AS DCs, comprising ∼1 in 5,000 PBMCs, had a nearly 5-fold enrichment in both *BCL11A* and *SPI1*-enriched populations (**Fig. 4g; Extended Data Fig. 4g**). Sub-clustering of the 269 cells from the sorted populations confirmed consistent expression of *AXL* and *SIGLEC6*, verifying their AS DC identity (**Fig. 4h**). As prior work reported two subclasses of AS DCs, including pDC-like and cDC-like cells, we hypothesized that our FlowFISH-sorted lineage-defining TFs may be associated with heterogeneity in this cell state. Indeed, differential gene expression analyses between the *SPI1*^+^ and *BCL11A*^+^ sorted populations identified key markers of the pDC-like subset (e.g.*, IL3RA, MZB1*) which are distinct from the cDC-like subset (e.g., *IFI30, ITGAX;* **Fig. 4i,j; Extended Data Table 4**). PERFF-seq differential analyses further identified new molecules that may play a functional role in these populations, including surface markers (*CD5, TLR9*), granzymes (*GZMB*), and cytokines (*IL1B*; **Fig. 4j; Extended Data Fig. 4h**).

Reanalysis of published Smart-seq2 data confirmed the co-expression of *BCL11A* and *SPI1* in these AS DC populations with marker genes previously identified for the cDC-like and pDC-like subpopulations^4^ (**Fig. 4k**). To further validate the identification of these TFs as AS DC subpopulation markers, we performed PERFF-seq targeting *IL3RA*, using an inclusive sort gate to include the *IL3RA*^low^ population (marked in our prior analysis with *SPI1^+^* FlowFISH; **Extended Data Fig. 4i**). From the 9,178 profiled cells, clustering analyses identified 95 AS DCs, and gene correlation analyses confirmed the separation of expression modules co-occurring with these two TFs (**Extended Data Fig. 4j,k**). Thus, analyses of our PERFF-seq datasets and the original Smart-seq2 profiles broadly corroborate the cDC-like and pDC-like subsets of AS DCs and nominate the lineage-defining *BCL11A* and *SPI1* TFs as candidate regulators of heterogeneity in this rare cell state.

Our analysis reveals key differences between protein and RNA-based sorting that affect the design of enrichment strategies. Specifically, *IL3RA* encodes CD123, an AS DC and pDC-specific surface marker, but our PERFF-seq profiles included many other T cell, B cell, and other myeloid populations, which is consistent with bulk mRNA profiles in PBMCs^18^ (**Extended Data Fig. 4l,m**). Further flow cytometry analyses confirmed distinct populations of CD123^+^/*IL3RA*^+^ populations (AS DCs and pDCs) as well as a distinguishable CD123^-^/*IL3RA*^+^ population (T cells, B cells, etc.; **Extended Data Fig. 4n**). These analyses collectively verify the sensitivity of PERFF-seq for identifying transcriptional heterogeneity in rare populations, and emphasize the power of nominating transcripts for enrichment or depletion based on transcriptional profiling data, compared to relying on classical conventions or surface protein marker expression alone.

### Enrichment and heterogeneity of rare nuclei revealed via PERFF-seq

Given the lack of well-described antibodies against nuclear protein markers, we asked whether PERFF-seq is compatible with the profiling of single nuclei from archived material, by isolating nuclei from fresh-frozen adult mouse brain cerebellum and FFPE-preserved human glioblastoma multiforme (GBM) tissue (**Fig. 5a**). Oligodendrocytes in the cerebellum are sparse (∼2% of total cells^20^) and are implicated in the cell-state-specific progression of multiple neurodegenerative disorders^21–23^, motivating us to profile them with the highly specific gene *Mobp*. For the human GBM samples (*n* = 2 donors), we selected three vasculature-associated genes, *DCN*^24^*, FN1*^25^, and *VWF*^26^, that have been individually implicated as biomarkers in glioblastoma pathogenesis or treatment response. Reanalysis of existing GBM FFPE data suggested that these three genes are principally expressed in a rare population (∼4.2%) of vascular-derived cells in primary GBM tumors (**Extended Data Fig. 5a,b**). Similar to our immune cell benchmarking (**Fig. 2**), downsampling analyses demonstrate that our upstream HCR-FlowFISH workflow has minimal, if any, effect on total UMI recovery compared to public 10x Genomics Flex profiles (**Fig. 5b,c; Methods; Extended Data Table 3**).

**Figure 5.**
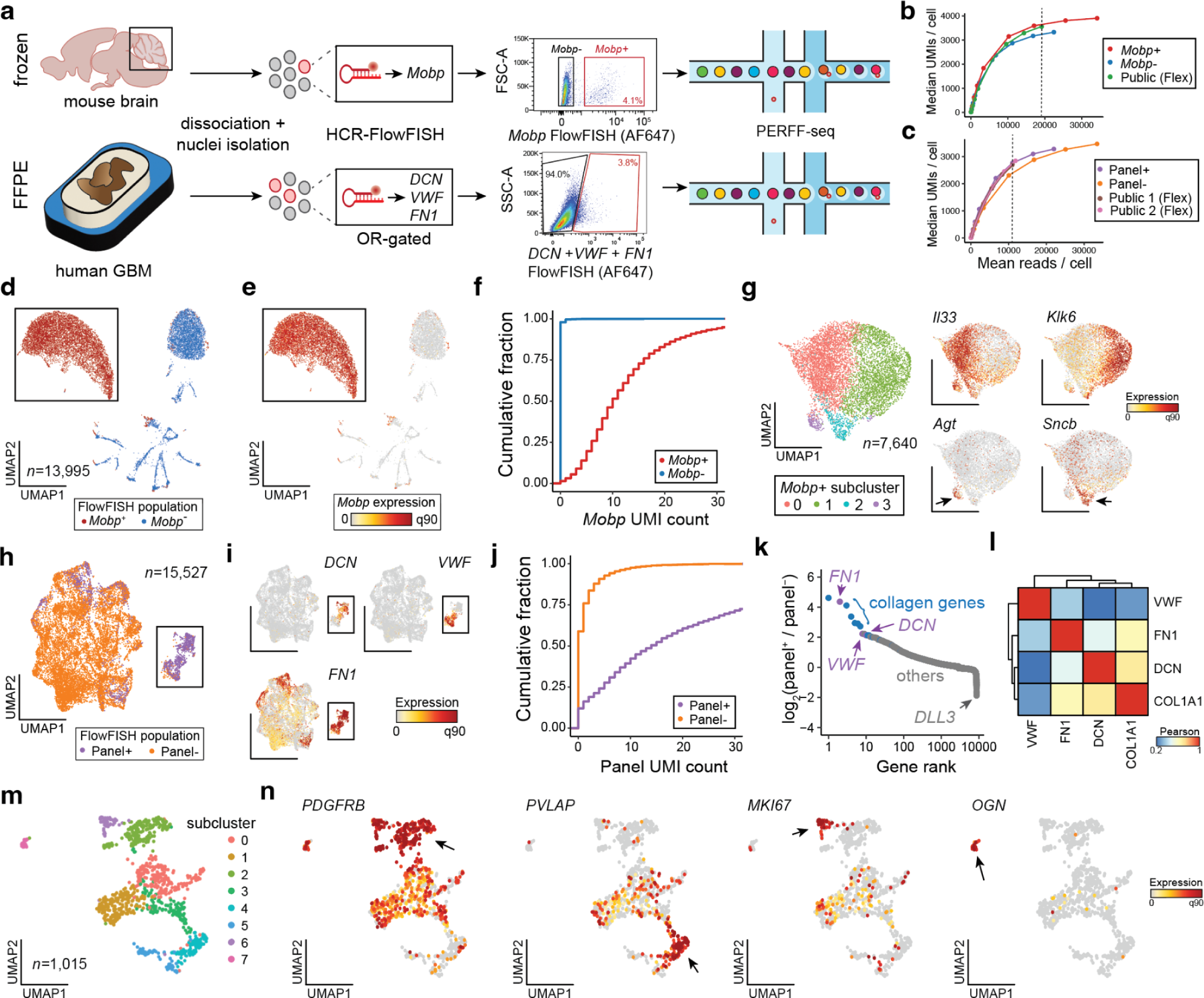
Study of rare nuclei from fresh and FFPE tissue. **(a)** Schematic of PERFF-seq single-nucleus experiments from frozen mouse brain tissue or FFPE human GBM tissue, showing HCR-FlowFISH staining and sorting strategy. **(b)** Downsampling analysis for library saturation and UMI benchmarking for the mouse brain nuclei. Dotted line represents the mean reads per cell for a final comparison (depth of lowest sample). **(c)** Same as (b) but for the human FFPE tissue sample. **(d)** UMAP embedding of the mouse brain nuclei FlowFISH enriched/depleted populations profiled with PERFF-seq. **(e)** Same as (d) but colored by *Mobp* marker gene expression. The boxed population was further subclustered. **(f)** Empirical cumulative distribution plot of raw UMI count for *Mobp* stratified by the captured PERFF-seq library. **(g)** Sub-clustering of the *Mobp*+ population. Arrows highlight top marker genes per cluster. **(h)** Reduced dimensionality representation of the human FFPE nuclei FlowFISH enriched/depleted populations profiled with PERFF-seq. **(i)** Same as (h) but colored by marker genes used in the FlowFISH panel. The boxed population was further subclustered. **(j)** Empirical cumulative distribution plot of total UMI count for the sum of the three genes enriched via FlowFISH, stratified by the captured PERFF-seq library. **(k)** Top differentially expressed genes between the two FFPE populations profiled with PERFF-seq. **(l)** Gene-gene correlations of relevant marker genes, including those used in the FlowFISH enrichment panel. **(m)** Sub-clustering of the Panel+ population with cluster states noted. **(n)** Top marker genes enriched in specific sub-clusters; arrows indicate critical populations where each gene is highly expressed.

Sorting and profiling the ∼4.1% of *Mobp*-expressing cells generated a nearly 50-fold enriched population (98.6% of cells *Mobp*^+^ cells have ≥1 *Mobp* UMI compared to 2.0% of the *Mobp*^-^ population) (**Fig. 5a,d,e,f**). Co-embedding of the *Mobp*^+^ and *Mobp*^-^ samples and annotation of canonical marker genes confirmed the expected cerebellum cell types, including granule cells (*Gabra6* and *Rbfox3*), interneurons (*Gad1*), and Bergmann glia (*Itih3*) that were detected in the *Mobp*^-^ library (**Extended Data Fig. 5c)**.

Subclustering on *Mobp*^+^ oligodendrocytes recovered four major populations with distinct marker profiles (**Extended Data Table 5)**. Cluster 0 (*n* = 3,805 nuclei) was characterized by *Il33* and *Ptgds*, known markers of terminally differentiated oligodendrocytes^27,28^ whereas cluster 1 (*n* = 3,194 nuclei) was characterized by *Klk6* and *S100B,* markers of maturing oligodendrocyte precursors that populate hindbrain structures and the spinal cord^29–31^ (**Fig. 5g, Extended Fig. 5d**). Clusters 2 and 3 were rare populations (*n* = 429 and *n* = 212 nuclei) defined by a mix of mature oligodendrocyte, oligodendrocyte precursor, and neuronal synapse markers (*Atp1b1, Scnb, Snap25* for cluster 2) in addition to *Agt* and *Aqp4*^20^ marking cluster 3, likely a low-frequency *Mobp*^+^ astrocyte population (**Fig. 5g, Extended Fig. 5d**). Though rare, the two subclusters likely represent distinct cell states—cluster 2 representing differentiating oligodendroglia that prune synapses^32^ and cluster 3 representing *Mobp^+^*astrocytes that have been described in cortical astrocyte populations^33^ but to our knowledge have not been identified in subcortical structures.

The PERFF-seq profiles from GBM enrichment exhibited a distinct cell cluster enriched primarily for cells from our vasculature OR-gated panel that express relevant marker genes (**Fig. 5h,i**). Not only did total UMI distributions in the panel clearly separate positive and negative populations (**Fig. 5j; Extended Data Fig. 5e**), but we detected a ∼23x increase in signal overall (mean panel^+^: 33.0 UMIs; mean panel^-^: 1.4 UMIs) where most background expression was driven by promiscuous expression of *FN1*, as expected (**Fig. 5i; Extended Data Fig. 5b**). Differential expression analyses confirmed genes enriched in our panel were among the top effects by fold-change with other collagen-associated genes similarly enriching within our vascular-enriched cells (**Fig. 5k**). Subclustering of the 1,015 vascular cells from our positive sort confirmed the two major populations, including *VWF*^+^ endothelial cells and *FN1*^+^/*DNC*^+^ mural cells (**Fig. 5l,m; Extended Data Fig. 5f**). While our identification of these major populations within the vasculature corroborates prior scRNA-seq analyses^34^, our high-quality PERFF-seq profiles allowed for further identification of 8 subclusters within the enriched vasculature (**Fig. 5m,n; Extended Data Fig. 5f; Extended Data Table 6**). These included an abundant population of *PVLAP*^+^ endothelial cells, a vascular marker of blood-brain barrier disruption^34^, proliferating cells marked by *MKI67*, and an ultra-rare population of mural cells expressing *OGN* that has not been well-defined in the human brain but is potentially linked to brain tumorigenesis^35^.

PERFF-seq logic-gating can thus be applied to enrich nuclei from rare populations upstream of single-cell profiling. Few, if any, high-quality antibodies against nuclear markers of rare cell types exist, emphasizing the need for such a programmable, RNA-based method for enrichment upstream of scRNA-seq for future Human Cell Atlas and disease profiling efforts.

## Discussion

PERFF-seq is a robust approach that can be used to enrich a variety of transcriptomic markers across distinct input material, including major immune cell populations (**Fig. 3**), very rare cell types (**Fig. 4**) and nuclei derived from fresh-frozen and FFPE material (**Fig. 5**). As solid tissue often requires or benefits from the isolation of nuclei rather than cells for profiling, the lack of high-quality intranuclear protein targets that meaningfully differentiate cell populations of interest poses a serious challenge. Furthermore, continued efforts to build cell atlases recover increasingly complex cell types defined by specific marker combinations, including in the mouse cerebellum^20^; thus, we envision a primary use of PERFF-seq to be enriching for specific populations defined by genes without the need for laborious genetic engineering, including via Cre-recombinases.

Though prior methods such as Probe-seq^36^ have coupled nucleic acid detection to bulk RNA sequencing, PERFF-seq is distinct in its ability to profile single-cell populations at high throughput (∼10^4^-10^5^ cells per capture). Another approach, FIND-seq^37^, sorts emulsion droplets to establish nucleic acid cytometry that similarly enriches rare cells based on specific transcripts. However, FIND-seq yields low-throughput (∼10^1^-10^3^ cells per capture) single-cell profiles and requires specialized instrumentation. We anticipate straightforward adoption of PERFF-seq by groups proficient in either RNA FISH or scRNA-seq, due to its combination of throughput and the commercial availability of reagents. While we showed experimental feasibility with up to three fluorophores in the same experiment (**Fig. 3**), we note that the HCR-FlowFISH kits contain 10 or more colors, allowing for further AND/NOT/OR logic-gating of populations before scRNA-seq profiling.

Though our dsDNase stripping effectively removes the HCR polymer, FISH probes likely remain bound to the target gene of interest. In our benchmarking with **Fig. 2**, we observed a ∼24% reduction in cells positive for *CD3E* and a two-fold log_2_ reduction in gene expression (**Extended Data Fig. 2d,e**). However, other targets such as *BCL11A*, *SPI1*, and *Mobp* showed robust expression as more than 90% of enriched cells were positive for the gene targeted by FISH. In addition, the Flex workflow requires ∼10^5^-10^6^ cells for cell pelleting upstream of droplet encapsulation, meaning that enriching for increasingly rare populations requires a concomitant abundance of starting material. We anticipate that these limitations can be overcome by using reagents for handling small numbers of cells^38^.

All droplet-based scRNA-seq workflows require substantial numbers of cells as input. The benefit of coupling HCR-FlowFISH FACS to scRNA-seq is that it enables distinct sorting strategies; rather than counting events on a cytometer, an inclusive PERFF-seq sorting strategy can be used that allows populations to be subsequently refined using transcriptomic profiles. For example, our application to sorting *BCL11A*^+^ cells allowed for detailed reanalyses of both AS DCs and B cells in the same experiment without pre-specifying the populations with distinct markers during sorting, as these populations could be readily separated by scRNA-seq analysis (**Fig. 4**). Though many surface markers that we evaluated such as *CD3E* and *MS4A1* (CD20) were consistent between surface protein and mRNA expression, others such as *IL3RA* (CD123) exhibited gene expression in lineages lacking surface protein expression. Similarly, though *BCL11A* is required for both B cell^39^ and pDC^40^ development, the higher mRNA expression of this TF in pDCs resulted in a substantial enrichment of this cell type and minimal enrichment of B cells in our PERFF-seq library.

Collectively, these vignettes motivate a careful data-driven exploration of appropriate marker genes using expression data rather than conventional knowledge derived from established FACS markers. Fortunately, such data-driven explorations are straightforward given the wealth of high-quality scRNA-seq profiles across many tissues, systems, and pathologies.

Ultimately, we anticipate that PERFF-seq will be particularly advantageous in settings where antibodies against marker proteins are either not available, not applicable, or poorly defined for a population of interest. In particular, as FlowFISH technologies have been established for viral gene expression^41^, ribosomal RNA content^42^, or long non-coding RNAs (**Fig. 2b**)^43^, our workflow provides unique access to the understudied populations defined by these markers. We envision an iterative process, whereby populations defined with existing large-scale scRNA-seq atlases are rationally enriched to study heterogeneity in new and rare populations, to refine cell atlases with ever greater definition.

---

**Extended Data Figure 1.**
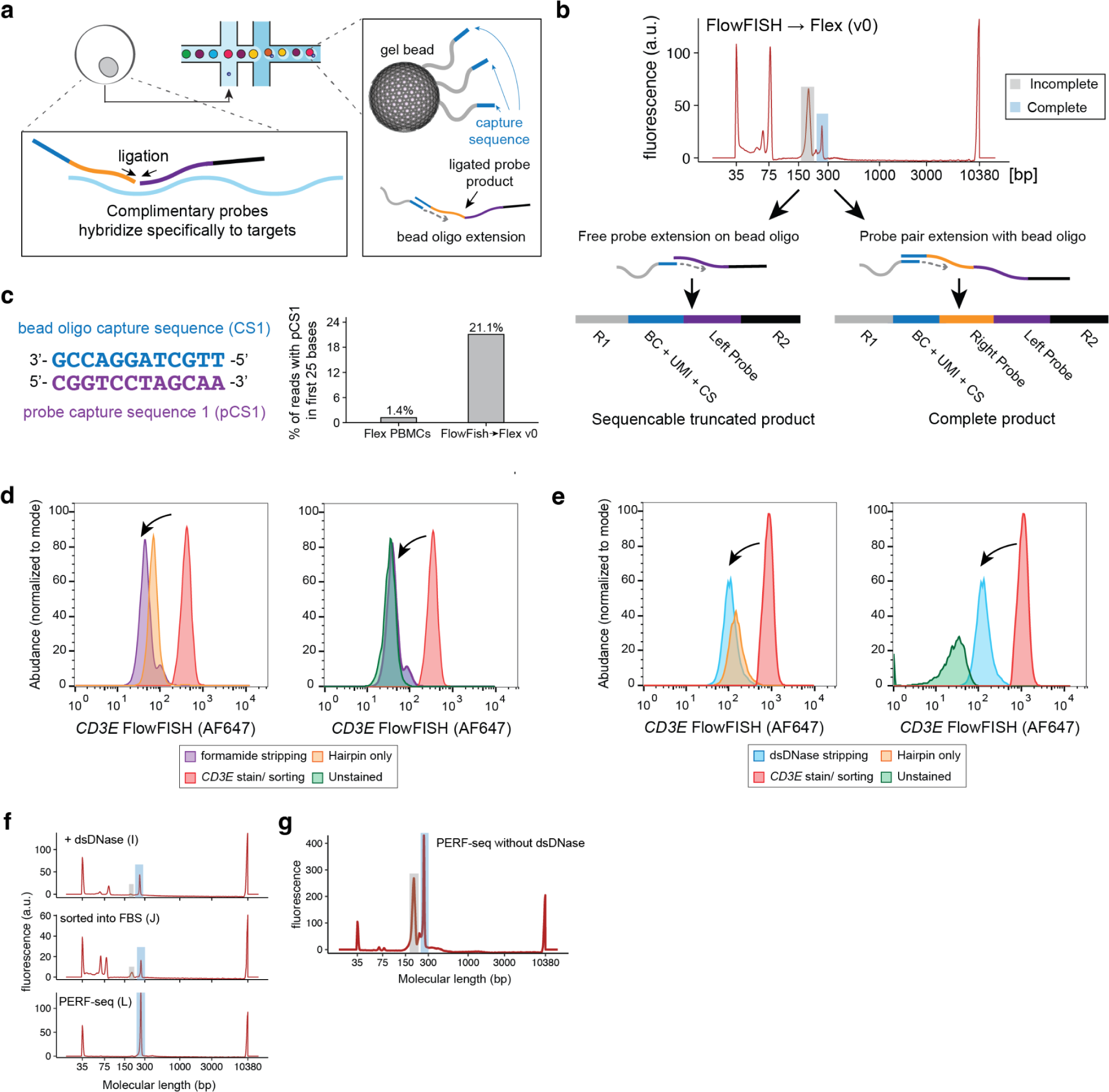
Analyses supporting PERFF-seq development. **(a)** Schematic overview of Flex workflow, including probe hybridization to transcript fragments in cells upstream of Chromium and bead oligo extension of the ligation product. (**b)** Representative Bioanalyzer (Agilent Technologies) trace outlining complete versus incomplete sequencing molecules. **(c)** Graphical summary of probe capture sequence (pCS1) - bead oligo capture sequence (left) and percent of reads with pCS1 detected in first 25 bases. **(d)** Comparison of FlowFISH signal using either unstained cells or the hairpin only in comparison to the sorted *CD3E* positive population and/or stripped via formamide. **(e)** Same as in (d) but using dsDNase for stripping. **(f)** Bioanalyzer traces for representative libraries from panels in Fig. 1, highlighting half- and fully-mapped probes. **(g)** Bioanalyzer traces of library preparation where the full PERFF-seq workflow was completed except for the omission of the dsDNAse stripping step.

**Extended Data Figure 2.**
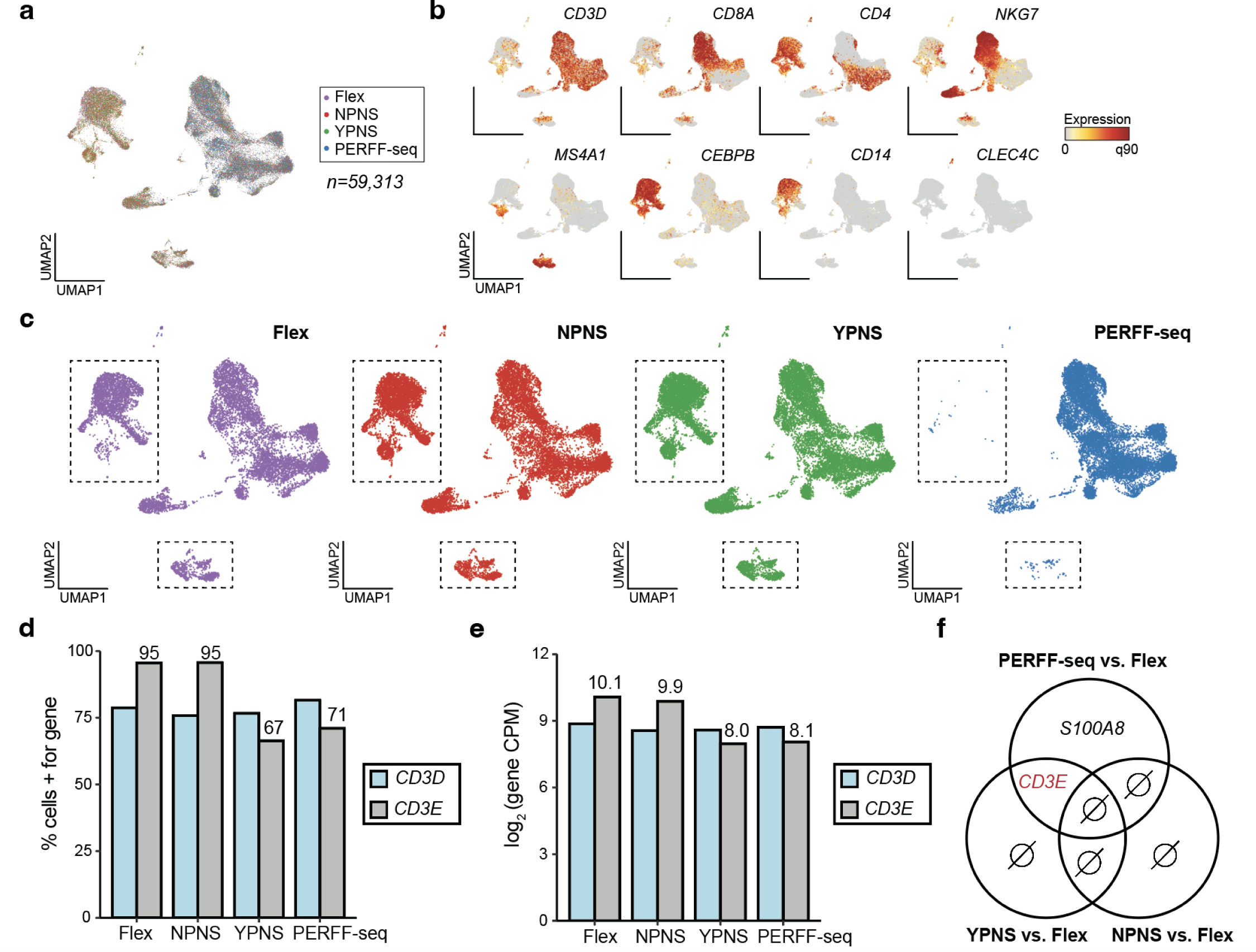
Supporting analyses of assay benchmarking. **(a)** Reduced dimensionality embedding of all cells in the four-plex benchmarking experiment. **(b)** Marker genes supporting annotation of key populations. **(c)** Same as (a) but stratified by library. Boxes indicate B cell and monocyte populations that are depleted from the PERFF-seq library. **(d)** Percent of T cells from each library with at least 1 UMI for *CD3D* or *CD3E*. **(e)** log2 counts per million (CPM) of *CD3D* and *CD3E* across different library conditions. **(f)** Differentially expressed genes between different facets of PERFF-seq compared to Flex. Two genes were differentially expressed, including *CD3E* in both the YPNS and PERFF-seq conditions. Ø means empty or no genes detected.

**Extended Data Figure 3.**
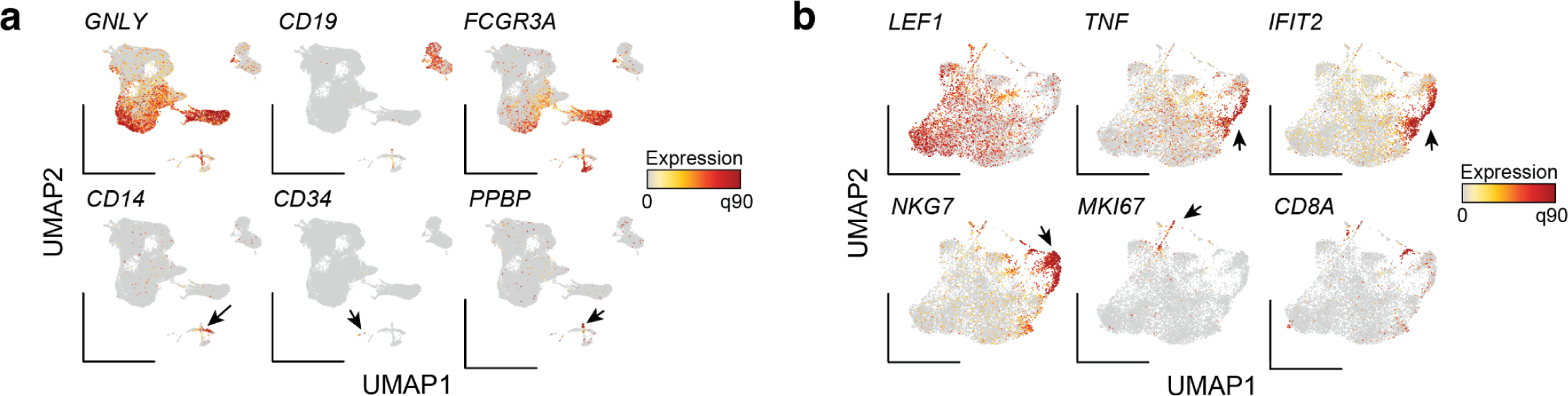
Supporting analyses of combinatorial PBMC cell states. **(a)** Additional marker genes for distinct populations from PBMC cell type analyses. Arrows indicate markers for rare populations expected from PBMC profiling. **(b)** Additional marker genes from *CD3E^+^ CD4^+^* sub clustering and rare population identification.

**Extended Data Figure 4.**
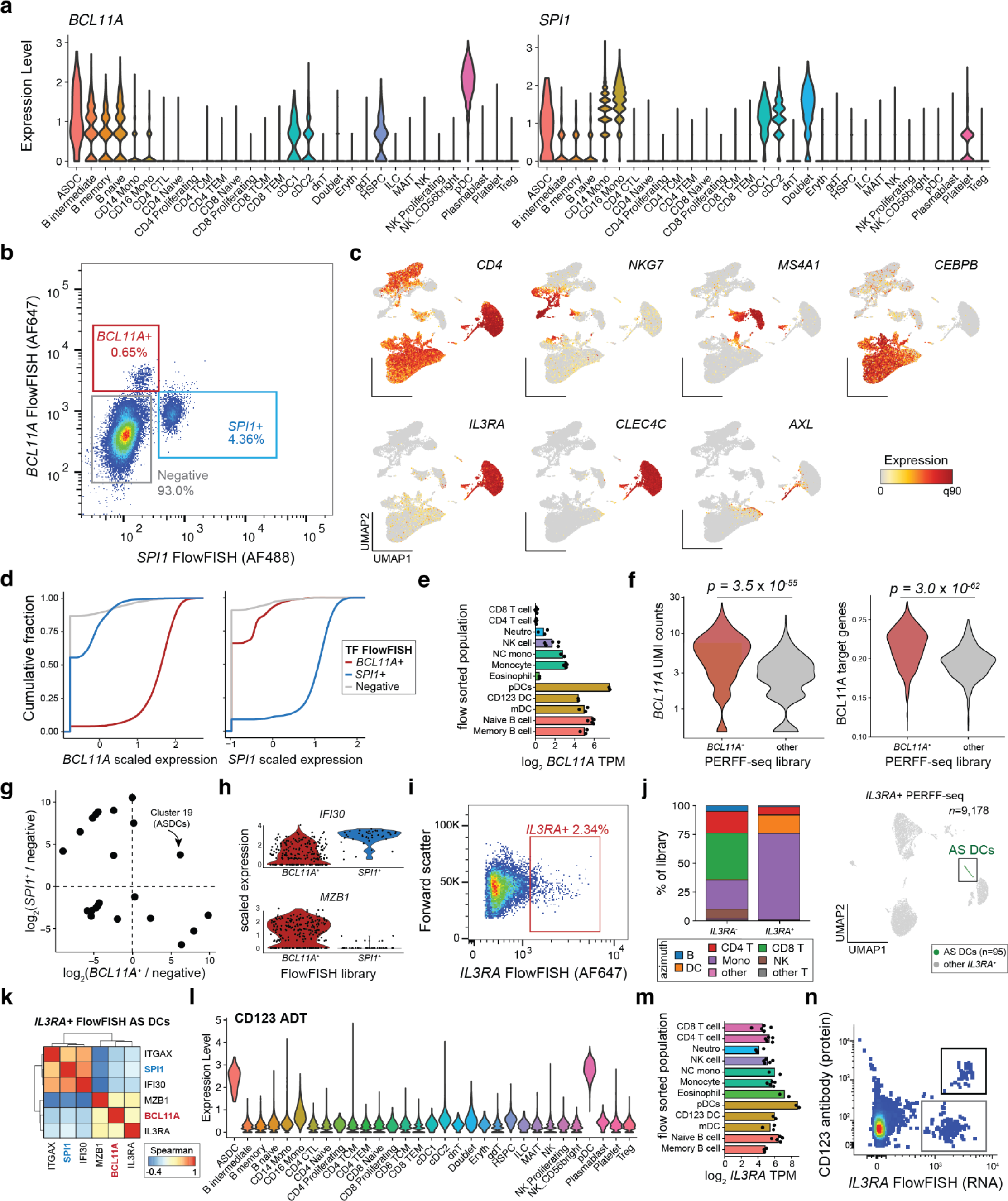
Supporting analyses of TF-based rare cell enrichments. **(a)** Azimuth Violin plots for *BCL11A* and *SPI1* RNA expression across well-annotated populations in peripheral blood mononuclear cell types. **(b)** Summary of FACS populations, including unsorted, *BCL11A^+^*, and *SPI1^+^*populations. **(c)** Additional marker genes supporting cell type annotations. **(d)** Empirical cumulative distribution plot of scaled expression of *BCL11A* (left) and *SPI1* (right) stratified by the captured PERFF-seq library. **(e)** Bulk RNA-seq of sorted populations of *BCL11A*^18^**. (f)** Comparison of B cells from *BCL11A*+ FlowFISH or negative/*SPI1^+^* populations for *BCL11A* expression or BCL11A target gene module scores. **(g)** Relative enrichment of each cell type in either the *BCL11A^+^* sort (x-axis) or *SPI1^+^*sort (y-axis) relative to the negative population using the raw Seurat cluster identities (compare to Fig. 4g). **(h)** Additional violin plots of marker genes, stratified by the FlowFISH library. All genes were significantly differentially expressed at a false discovery rate (FDR) < 0.01. **(i)** Summary of *IL3RA^+^* FACS sort and population characterized with PERFF-seq. **(j)** Left: proportion of cell types from the Azimuth L1 reference for *IL3RA*+ and - PERFF-seq libraries. Right: Reduced dimensionality representation of *IL3RA^+^* PERFF-seq library, highlighting profiled AS DCs. **(k)** Gene-gene correlations of all AS DCs from the *IL3RA^+^* sort. Genes match those in Fig. 4k. **(l)** Summary of CD123 expression from antibody-derived tags (ADT) of PBMC CITE-seq. **(m)** Bulk RNA-seq of sorted populations of *IL3RA*^18^. **(n)** Flow cytometry analysis of PBMCs co-stained with *IL3RA* mRNA (via FlowFISH) and CD123 protein (encoded by *IL3RA*). Double and single positive populations are indicated in the boxes.

**Extended Data Figure 5.**
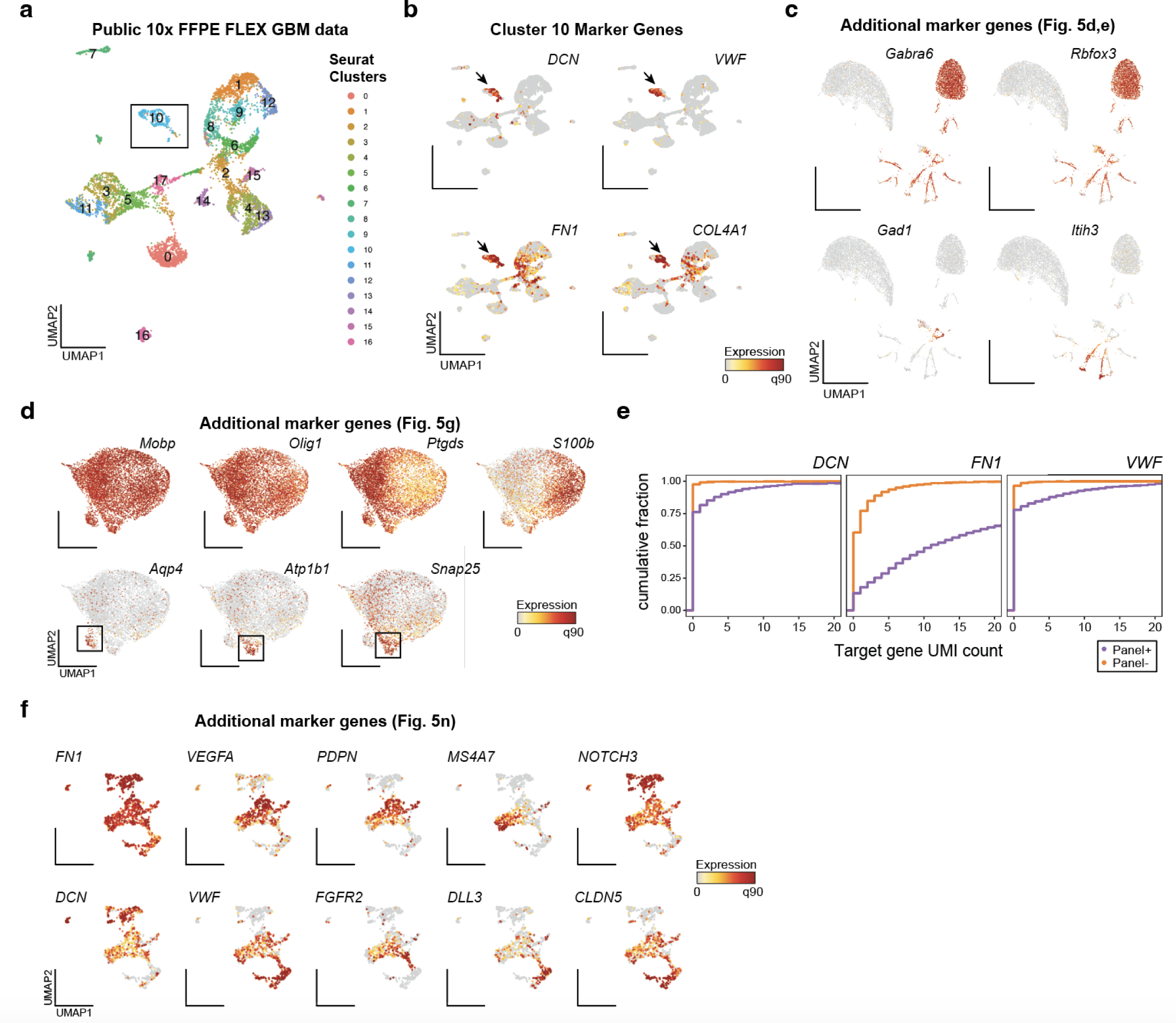
Supporting analyses of nuclei enrichment from fresh and fixed tissues. **(a)** Reduced dimensionality representation of public GBM FFPE Flex data showing 17 clusters. **(b)** Annotation of marker genes for cluster 10, the population highlighted by the arrow. **(c)** Supporting marker genes annotating other subpopulations from the PERFF-seq experiment, including the primary cluster of granule cells. **(d)** Additional marker genes from *Mobp^+^* cells were profiled with PERFF-seq. Atp-associated genes supporting rare subclsuters are noted in the boxes as well as marker genes highly expressed in all cells. **(e)** Empirical cumulative distribution plot of the raw UMI counts for each of the three genes enriched via FlowFISH, stratified by the captured PERFF-seq library. **(f)** Additional marker genes showing heterogeneity defining subclusters of endothelial cells and pericytes.

## Extended Data Tables

**ED Table 1. Summary of probes used in the manuscript.**

**ED Table 2. Summary of Flex libraries profiled.**

**ED Table 3. Statistics comparing PERFF-seq libraries for benchmarking.**

**ED Table 4. Differentially expressed genes between AS DC subpopulations.**

**ED Table 5. Marker genes in mouse brain oligodendrocytes.**

**ED Table 6. Marker genes in human endothelial cells from GBM FFPE tissue.**

## Methods

### PERFF-seq method and development

A full protocol for executing the combined HCR-FlowFISH and 10x Genomics Flex profiling steps is available on protocols.io (dx.doi.org/10.17504/protocols.io.14egn3k6ql5d/v1 dx.doi.org/10.17504/protocols.io.8epv5x8r4g1b/v1) and in the following methods section. In brief, based on our optimizations in **Fig. 1**, we emphasize a few critical aspects of the method development. First, enriching for populations via HCR-FlowFISH proceeds with minimal modifications to the protocol except for the use of RNase-inhibitor BSA buffer to preserve RNA quality. After the enriched populations are isolated, a dsDNase step is used to degrade the HCR-polymer, which we found is critical for 10x Genomics library preparation. After dsDNase digestion, the inclusion of the 10x Flex WTA probes proceeds as is standard for the workflow before droplet encapsulation on the 10x Genomics Chromium platform. Downstream amplification and sequencing follow the standard Flex guidelines with no modifications. Together, PERFF-seq leverages the quality-controlled aspects of both workflows with minimal modifications, but we emphasize that these modifications (dsDNase and RNase-free BSA) can severely limit data quality if left unaccounted. Complete details for each step are available from the manufacturer as well as the protocols.io link accompanying this manuscript.

### Quality control metrics and optimization

From both Bioanalyzer traces and CellRanger quality control summaries, we observed the presence of half-ligated probes was a correlate for overall library quality (**Fig. 1; Extended Data Fig. 1**). Our interpretation of the half-mapped probes is that conditions for Flex are incompatible with the ligation required for the pairs of gene probes for RNA detection (**Extended Data Fig. 1**). As a consequence, many left probes become barcoded, resulting in a product that is ∼70 bases smaller than the expected Flex barcode product (**Extended Data Fig. 1b**). To confirm this, we examined one million reads from either of our initial libraries for the presence of the probe capture sequence (“CGGTCCTAGCAA”) in the read position where the left probe sequence is contained on R2 (between positions 1 and 25), which resulted in ∼21.1% of reads containing a perfect match to probe capture sequence (**Extended Data Fig. 1c**). From our remaining optimization experiments, our interpretation of these data is that the presence of the HCR-Polymer (or formamide) are sufficient to disrupt the essential probe ligation for high-quality flex data. In this sense, the removal of the polymer after sorting via dsDNase removes the polymer that otherwise inhibits ligation.

### Immune cell experiments

Protocol development and optimization were performed on cryopreserved peripheral blood mononuclear cells (PBMCs) sourced from ATCC and AllCells. Vials were thawed and viability exceeded 90% for all samples. PBMCs were used as the primary input for developing the assay in **Fig. 1** due to the ease of material availability and well-defined heterogeneity for *MS4A1* and *CD3E*. For applications in **Fig. 2-4**, the same vials were used but enriched for specific markers as indicated in the experimental overview schematics (**Fig. 2c, 3a, 4a**). All experiments started with ∼10M cells, except for the TF sort experiment (**Fig. 4**), which began with ∼25M cells to yield ample cell numbers for downstream profiling given the rare *BCL11A* population that was sorted.

### Cell Fixation and permeabilization

Fixation and permeabilization were performed as described in “HCR RNA flow cytometry protocol for mammalian cells in suspension” provided by molecular instruments. Briefly, cells were thawed and fixed in 4% paraformaldehyde solution (4% paraformaldehyde in 1x PBS and 0.1% Tween 20) at room temperature for 1 hour at a concentration of 1 million cells per mL (1M/mL). After fixation, cells were centrifuged at 350 xg for 5 minutes and resuspended in a PBST solution (1x PBS and 0.1% Tween 20) at 1M/mL. This step was repeated once for a total of 2 washes. After washing, cells were permeabilized with ice-cold 70% EtOH overnight at 1M/mL. After permeabilization, cells were centrifuged and resuspended with PBST solution at 1M/mL twice.

### HCR FlowFISH

Probes for RNA targets of interest and complementary hairpins were purchased from Molecular Instruments at the highest number of probe pairs available for genes of interest.

Most steps were performed as described in the “HCR RNA flow cytometry protocol for mammalian cells in suspension” provided by molecular instruments and Reilly et al^13^ with the following adjustments: First, all centrifugation was performed at 850 xg for 5 minutes unless otherwise noted. Second, low-binding plasticware tubes and RNase-free molecular-grade reagents were utilized when possible. Third, during the detection stage, we found that an optimal signal-to-noise ratio was achieved during fluorescence detection at 16nM probe concentration per 500,000 - 1 million cells (8uL of probe stock per sample). Finally, 37^0^C incubations were performed in a heated lid thermomixer with gentle shaking.

#### HCR FlowFISH Detection Stage

Permeabilized cells/nuclei were resuspended in pre-warmed 400uL of hybridization buffer (molecular instruments) per 500,000 to 1M cells. Cells/nuclei were incubated for 30 minutes at 37C, 300rpm in a heated lid thermomixer. The probe solution was prepared by mixing 8uL of 1uM probe stock and hybridization buffer for a final 100uL volume per sample. Probe solution was added to each sample for a final probe concentration of 16nM and cells were incubated at 37C for 16-24 hours. To pellet cells/nuclei, 500uL of SSCT solution (5X SSC, 0.1% Tween 20) was added to each sample and centrifuged at 850xg for 15 minutes. Cells/nuclei were resuspended in 500uL of prewarmed probe wash buffer (molecular instruments), incubated for 10 minutes at 37C, and subsequently pelleted at 850xg for 5 minutes. This step was repeated 3 more times for a total of 4 washes. Then, cells/nuclei were resuspended in 500uL of SSCT solution and incubated at room temperature for 5 minutes.

#### HCR FlowFISH Amplification Stage

Cells/nuclei were centrifuged and resuspended in 150uL of amplification buffer and incubated at room temperature for 30 minutes. In the meantime, 5uL of 3uM h1 and h2 hairpin stock was aliquoted for each probe set and snap cooled by performing a heat shock at 950C for 90 seconds and cooling in the dark at room temperature for 30 minutes. To prepare the hairpin solution, snap-cooled h1 and h2 hairpins were mixed with an amplification buffer to make a final volume of 100uL per sample. The hairpin solution was added to appropriate samples for the final hairpin concentration of 60nM. Cells/nuclei were incubated at room temperature for 16-24 hours. (This time can be reduced to 4 hours). After incubating, samples were washed 6x with 500uL of SSCT for each sample.

#### Co-staining with antibodies

PBMCs were thawed and stained with anti-CD123 antibody (Biolegend S18016F; **Extended Data Fig. 4n**). Antibody staining was immediately followed by fixation/permeabilization and HCRFlowFISH as described above. We note that surface antibodies conjugated to synthetic dyes result in the most robust signal when used in conjunction with the HCR-FlowFISH protocol.

### Sample Enrichment using FACS

Samples were resuspended in sorting and collection buffer (1x PBS, 5% BSA - Gibco #15260037, 0.13U/uL RNAse inhibitor - Millipore Sigma #3335399001) and filtered through a 35um strainer. Cells were kept in dark and on ice until sorting. Collection tubes were prepared with 300uL of collection buffer. Cells were sorted using the BD FACSAria III or FACSymphony S6 with 70um nozzle for cells and 85um nozzle for nuclei. Representative gating is shown where appropriate. For multiplexing experiments, compensation was performed with single-color controls.

### HCR Polymer Disassembly

Sorted cells were pelleted and resuspended in 275uL of 1x dsDNase buffer (Thermofisher #EN0771) and incubated for 15 minutes after which 25uL of dsDNase enzyme (Thermofisher #EN0771) was added and the sample was incubated at 37^0^C for 2 hours. After incubation, 3uL of 1M DTT was added to the sample to quench dsDNAse activity and incubated at 55^0^C for 5 minutes for heat inactivation. Samples were pelleted at 850xg for 5 minutes, resuspended in 500uL of pre-warmed wash buffer, and incubated for 10 minutes. This step was repeated once for a total of 2 washes. Samples were then resuspended in 500uL of SSCT buffer and incubated for 5 minutes. Samples were centrifuged and resuspended in 1mL of 0.5x PBS and 0.02% BSA (Thermofisher #AM2616) and 0.2u/uL RNAse inhibitor (Millipore Sigma #3335399001). Polymer Disassembly was assessed by measurement of fluorescence intensity on the BD FACSAria III.

### 10x Genomics Flex

Preparation of all 10x Genomics Flex libraries were prepared using the manufacturer’s instructions as all modifications for PERFF-seq happen upstream of Flex probe hybridization. All libraries were sequenced on an Illumina Nextseq 550, Novaseq 6000, Novaseq X, or an Element AVITI with standard dual-indexing and demultiplexing. Raw .bcl files were processed using CellRanger v7.2, and the resulting .fastq files were quantified for the human and mouse probe set to version 1.0.1 using default parameters for the CellRanger pipeline.

### Cell line mixing and benchmarking

The XY Burkitt’s lymphoma cell line (Rajis) and the XX Chronic Myelogenous Leukemia cell line K562 cells were obtained from ATCC. Raji cells were cultured in RPMI-1640, while K562 cells were cultured in Iscove’s Modified Dulbecco’s Medium (IMDM), both supplemented with 10% fetal bovine serum and 1% Penicillin/Streptomycin. To benchmark the recovery of populations, cells were washed with PBS, centrifuged, and counted. Raji cells were subsequently mixed with decreasing 10-fold dilutions of K562 cells. The mixed cells were then fixed and permeabilized as described in the prior sections followed by the HCRFlowFISH protocol using *XIST* RNA probes for detection, and Alexa Fluor 647-conjugated hairpins for amplification. FACS analyses were conducted on ThermoFisher Attune NxT Flow Cytometer.

### Mouse Tissue Sourcing

Mouse brain tissue was sourced from Zyagen Inc. as a fresh frozen whole brain stored in OCT. Upon receipt, tissue was stored at −80C. Dissection was performed in a cryotome and immediately processed for nuclei processing.

### Mouse brain dissociation and profiling

Mouse Brain Dissociation was performed as described by 10x genomics in the tissue fixation and dissociation for Chromium Fixed RNA Profiling. Briefly, fresh frozen mouse cerebellum was weighed and fixed in 4% paraformaldehyde solution for 2 hours at 2mL per 25mg of tissue with periodic agitation. Then, the tissue was centrifuged and re-suspended in 1x PBS twice. Washed tissue was resuspended in ice-cold 70% ethanol at 2mL per 25mg of tissue and incubated overnight at 4C. After incubation, the tissue was centrifuged and resuspended with 1x PBS twice. Tissue was then resuspended in 2mL dissociation buffer (160uL LiberaseTL enzyme - Millipore Sigma #5401020001 + RPMI) using the gentleMACS OctoDissociator with heaters (Miltenyi Biotec # 130-096-427) for 30 minutes at 50rpm. Nuclei were washed with 1x PBS + 0.02% BSA (Thermofisher #AM2616) + 0.2u/uL RNAse inhibitor and stained 1ul/ml DAPI(Thermofisher #62248) for 10 minutes. To remove excess debris, DAPI^+^ nuclei singlets were sorted using FACS before processing by HCRFlowFISH. Nuclei were either stored for future use according to 10x Genomics Recommendation or proceeded directly into HCRFlowFISH.

### GBM FFPE dissociation

FFPE samples were preprocessed on a prototype S2 Singulator system. The sample was automatically processed in a NIC+ cartridge (S2 Genomics #100-215-389) by three 15 minute deparaffinization steps (CitriSolv, VWR), rehydrated by successive 1 mL washes of 100%, 100%, 70%, 50%, and 30% ethanol, followed by 2 washes of PBS. The sample was then spun at 1,000g for 3 min and resuspended in 0.5 mL Nuclei Isolation Reagent (NIR, S2 Genomics, #100-063-396) with 0.1 ul/uL RNase inhibitor (Protector, Millipore Sigma, #3335399001); all subsequent solutions had RNase inhibitor. The sample was dissociated to single nuclei in a second NIC+ cartridge with 2 mL of NIR for 10 min followed by a 2 mL wash with Nuclei Storage Reagent (NSR, S2 Genomics, #100-063-405). The single nuclei suspension was spun 500g for 5 min, resuspended in NSR, and counted.

### Bioinformatics analyses overview

All bioinformatics analyses were conducted using standard output files from the execution of CellRanger to sequencing data of the Flex libraries. Downstream analyses, including cell filtering, marker gene analyses, and visualization, were performed using Seurat v4^19^. In brief, cells were identified via a combination of passing the CellRanger knee plot as well as meeting minimum quality control standards, including at least 1,000 UMIs detected, 500 genes detected, and no more than 5% mitochondrial RNA abundance, which are standard thresholds for scRNA-seq analyses. For all sub-clustering analyses (**Fig. 3g, 4h, 5g, 5m**), we required cells to be both present in the enriched PERFF-seq library as well as belonging to the Seurat cluster associated with the majority of the population. All differential expression and marker gene analyses were performed using the FindMarkers functionality in Seurat^19^. All custom code to reproduce all custom downstream analyses, including intermediate data files is available as part of an online repository.

### Benchmarking analyses

To examine the loss of data quality in PERFF-seq compared to analogous Flex libraries, all comparisons were made against gold-standard data generated and released by 10x Genomics. Saturation curves were drawn by downsampling the total reads in the library to 0.1%, 1%, 2.5%, 5%, 10%, 30%, 50%, 75%, and 100% of the total sequencing depth. Downsampling proceeded via the sample() function in R on the per-molecule vectors encoded in the _sample_molecule_info.h5 file from the CellRanger processing. To compare median per-cell UMI and gene counts, we selected the read depth of the lowest library in the comparison and downsampled every other library to compare relevant statistics. Thus, these analyses are robust to differences in sequencing depth, UMI collapsing, and barcode correction (which occur within the CellRanger processing steps upstream of the h5 output).

To assess the impact of the PERFF-seq workflow compared to Flex, the *CD3E* targeted transcript was analyzed alongside *CD3D* (**Extended Data Fig. 2d-f**). For these panels, we subsetted the analyzed cells to the distinct clusters of T cells from the Seurat embedding. Differential gene expression analyses were conducted using the FindMarkers function and requiring a log2FC of 0.5 for inclusion in the Venn diagram, resulting in only two genes, *CD3E* and *S100A8* across the three pairs of comparisons. These results indicate that the overall transcripts of T cells were not changed between the potential conditions, though there is a ∼4x reduction in *CD3E* expression where ∼70% of T cells had detectable levels of *CD3E* (compared to 95%) from Flex alone. As other genes (e.g., *BCL11A*, *SPI1*, *Mobp*) had ∼90% of cells sorted via FlowFISH have detectable expression from the Flex library, we suggest that each gene may behave differently, likely as a function of overall gene length and whether the Flex and HCR-FlowFISH probes overlap at the same locus on the mRNA. Thus, special consideration is required for the specific genes used in the FlowFISH panel, but the overall transcripts of the cells should remain stable.

### Immune cell analyses

To benchmark the enrichment efficiency (**Fig 2e**; **Fig. 3d**), we utilized three distinct methods. First, reference projections with azimuth to a gold-standard PBMC atlas were utilized, noting the differences in chemistries between the reference and projections (reference: reverse-transcription-based; here: probe-based Flex chemistry). Qualitatively, the cell type assignments from azimuth were sensible but projection onto the two-dimensional space often failed to cover the breadth of the reference, which we attribute to differences in the fundamental sequencing chemistry. As a second reference-based method, we annotated genes using the default celldex^44^ workflow for human immune cells. For the logic-gated classification (**Fig. 3**), we partitioned the output from classification as “B cells” for *MS4A1*^+^, “CD4+ T cells” for *CD4*^+^*,CD3E*^+^ cells, “CD8+ T cells” and “T cells” for the *CD4-,CD3E*^+^ population, and all other labels as the negative population. Finally, we individually clustered genes and defined cell type annotations based on standard practice for the presence or absence of individual marker genes. The proportions annotated as accurate classification represent the total number of high-quality cells (*n*>10,000 per comparison) and were consistent between different classification methods, verifying the specificity of our enrichment via sorting strategy and preservation of transcriptomes for downstream analyses. For comparisons with other RNA-seq datasets, normalized data from flow-sorted bulk^18^ populations and single-cell annotations^19^. A collection of 2,674 target genes of BCL11A was downloaded from Harmonizome 3.0^45^ using the standard AddModuleScore functionality in Seurat^19^.

### Nuclei analyses

Mouse brain analyses, including clustering and sub-clustering analyses, proceeded as described above. The selection of plotted marker genes followed the cerebellum atlas that defined oligodendrocyte subtype markers without any prior enrichment^20^. To compare PERFF-seq performance against frozen nuclei, we downloaded a public four-plex FLEX library and selected the dissociated eye nuclei as the closest anatomical tissue to our profiled tissue, noting these are an imperfect yet useful comparison (**Fig. 5b**).

For existing GBM FFPE Flex data, the counts matrix was downloaded from the datasets hosted on the 10x Genomics website. The two GBM samples were run as a 4-plex in-line barcode multiplexing with another tumor type (colorectal) which was discarded during pre-processing. These processed data were used both in defining endothelial/mural cell markers (**Extended Data Fig. 5a,b**) as well as in the downsampling performance analysis (**Fig. 5c**). Selection of marker genes was based on prior profiles of endothelial cells from GBM cells^34^.

## Acknowledgments

We are grateful to the Satpathy Lab, Lareau Lab, Gladstone Flow Cytometry Core, and Single-cell Analytics Innovation Lab members for helpful discussions. We acknowledge a helpful blog post from 10x Genomics describing the singlet unligated probe set. We thank T. Nawy for helpful feedback on the manuscript and method. We extend our gratitude to Nathan Pereira and Stevan Jovanovich of S2 Genomics for their assistance and support in the preparation of nuclei from FFPE Tissue. This work was supported by NIH grants R00HG012579 (C.A.L.) and UM1HG012076 (A.T.S.; L.S.L.). A.T.S. is supported by the Burroughs Wellcome Fund Career Award for Medical Scientists, the Parker Institute for Cancer Immunotherapy, a Pew-Stewart Scholars for Cancer Research Award, a Cancer Research Institute Lloyd J. Old STAR Award, a Scholar Award from the American Society of Hematology, and a Baxter Foundation Faculty Scholar Award. YHH is supported by a PhD fellowship from the Hector Fellow Academy. LSL is supported by the Hector Fellow Academy, the Paul Ehrlich Foundation (LSL), the EMBO Young Investigator Programme (LSL), an Emmy Noether fellowship (LU 2336/2-1) and grants by the German Research Foundation (DFG, LU 2336/3-1, LU 2336/6-1, STA 1586/5-1, TRR241, SFB1588, Heinz Maier-Leibnitz Award). SAIL is supported by the Alan and Sandra Gerry Metastasis and Tumor Ecosystems Center (GMTEC) at MSK.

## Author contributions

T.A., R.R.S., M.T., R.C., A.T.S., and C.A.L., conceived and designed this work. T.A. and R.R.S. led assay development with input from A.T.S., and C.A.L. T.A., R.R.S., M.T., B.N.N., Y.-H.H., S.H., and C.S. performed experiments. K.K.H.Y., Z.A.-M., V.T., and L.S.L. aided in the interpretation of data analyses. R.R.S., R.C., A.T.S., and C.A.L. supervised the work. C.A.L. led bioinformatics analyses with input from T.A. and R.R.S. T.A. led the development of the protocol with input from R.R.S. and M.T. T.A., R.R.S., and C.A.L. drafted the manuscript with input from other authors.

## Code and Data Availability

Sequencing data associated with this work is available at GEO accession **GSE262355**. Custom code and intermediate data files to reproduce all analyses supporting this manuscript are available at https://github.com/clareaulab/perffseq_reproducibility. A full step-by-step protocol for PERFF-seq is available on protocols.io (dx.doi.org/10.17504/protocols.io.14egn3k6ql5d/v1, dx.doi.org/10.17504/protocols.io.8epv5x8r4g1b/v1).

